# Spike-history gating of plateau potentials enables closed-loop rewriting of CA1 representations

**DOI:** 10.64898/2026.09.15.751647

**Authors:** Anno C. Kurth, Toshitake Asabuki

## Abstract

Behavioral time scale synaptic plasticity (BTSP) allows hippocampal CA1 neurons to form or shift place fields after a single plasticity-inducing plateau potential, but what determines when individual neurons become eligible for another plasticity event remains unclear. Here we propose that recent somatic spike history provides a cell-specific gate for plateau initiation, making neurons sensitive to rising activity after relative silence while suppressing repeated plasticity during sustained firing. In a two-compartment spiking model of CA1, this gate interacts with entorhinal input to create a feedback loop in which existing representations bias subsequent plateau locations. The resulting dynamics produce gradual place-field drift, reward-associated overrepresentation, recovery of environment-specific representations after remapping, and cross-day reinstatement. Our model further predicts that stronger residual activity biases subsequent plateaus toward previous field locations, a relationship we identify by reanalyzing experimental data. Thus, the expression of neural representations can help determine when and where they are subsequently rewritten.

## 1 Introduction

Many classical forms of synaptic plasticity rely on precisely timed and repeated coactivity of the involved neurons [1, 2, 3]. Behavioral time scale synaptic plasticity (BTSP) provides a strikingly different form of plasticity in hippocampal CA1. A single dendritic plateau potential (PP) can trigger large synaptic changes over seconds-long windows and form or shift place fields within single trials [4, 5]. BTSP-like mechanisms have since been reported in hippocampal CA3 and visual cortex [6, 7, 8], suggesting a broader role for event-triggered plasticity, and computational implications have been explored for one-shot learning and the credit assignment problem [9, 10, 11]. Unlike plasticity that accumulates gradually through repeated coincident pre- and postsynaptic activity, BTSP depends on the time between the onset of the plateau and the presynaptic activity. This poses a fundamental problem for the post-synaptic neuron: how to control the timing of the plasticity-inducing events?

Experimental work highlights inputs from entorhinal cortex layer 3 (EC3) as the main driver of plateau potential initiation [12, 13, 14]. Manipulation of EC3 activity can regulate place-field formation [14], and reward related fluctuations can lead to place-field overrepresentation [13]. This motivated the proposal that EC3 provides a target signal that instructs where and how strongly BTSP is favored [13, 15]. Yet individual EC3 axons show broad spatial tuning and substantial trial-to-trial variability, with only a minority (around 19%) showing highly reproducible spatial firing, suggesting a population rather than single-neuron level control via EC3 input [16].

At the same time, place field changes in individual CA1 neurons over time are often smooth rather than discontinuous place field translocations [17, 18]. Stochastic BTSP induction driven by neural spiking can account for such gradual drift [18], but firing level alone does not explain how an individual neuron distinguishes activity that should permit another plasticity event from sustained firing at an already expressed field.

Here, we propose that recent somatic spiking history provides a means for this cell-specific control. Building on the coupling between somatic firing, back-propagating action potentials, and dendritic calcium electrogenesis [19, 20], and on the recently discovered spike-rate accelerometer mechanism [21, 22] we suggest that plateau initiation becomes possible when spiking activity increases rapidly after a period of low activity, while sustained high activity suppresses further events. Somatic history thus gates when an individual neuron is available for a plateau potential, while EC3 and local inhibition control whether a plateau is generated once the gate is open. In a two-compartment view of CA1 pyramidal cells, this interaction closes a feedback loop in which existing place fields bias subsequent plateau potentials, which in turn reshape future place fields. This mechanism also addresses a sensitivity-stability dilemma for synaptic plasticity in place cells: the gate remains sensitive to the onset of weakly or partially formed place-field activity, while history dependence together with post-plateau refractoriness limits unnecessary plasticity at strongly expressed fields.

We tested this idea in a stylized computational model that produces gradual place field drift, allows reward-related EC3 modulation to bias plasticity, preserves environment-specific representations across remapping, and enables weak residual traces to guide cross-day place field reinstatement. Importantly, residual activity from previous fields biases subsequent PPs toward their former locations, a relationship we also identify by reanalyzing recent experimental data. Our results shed light on a reciprocal relationship between neural activity and plasticity, where the expression of representations helps to determine the timing and location of their own future modification.

## 2 Results

### 2.1 Spike-History-Dependent Plateau Gating

EC3 has been proposed to provide an instructive or target-like signal for plateau-dependent plasticity in CA1 [12, 13, 14] (Figure 1A, left). Such input biases where and how strongly plasticity is favored. This leaves open the question what determines whether an individual CA1 neuron is currently available for another plasticity event.

**Figure 1:**
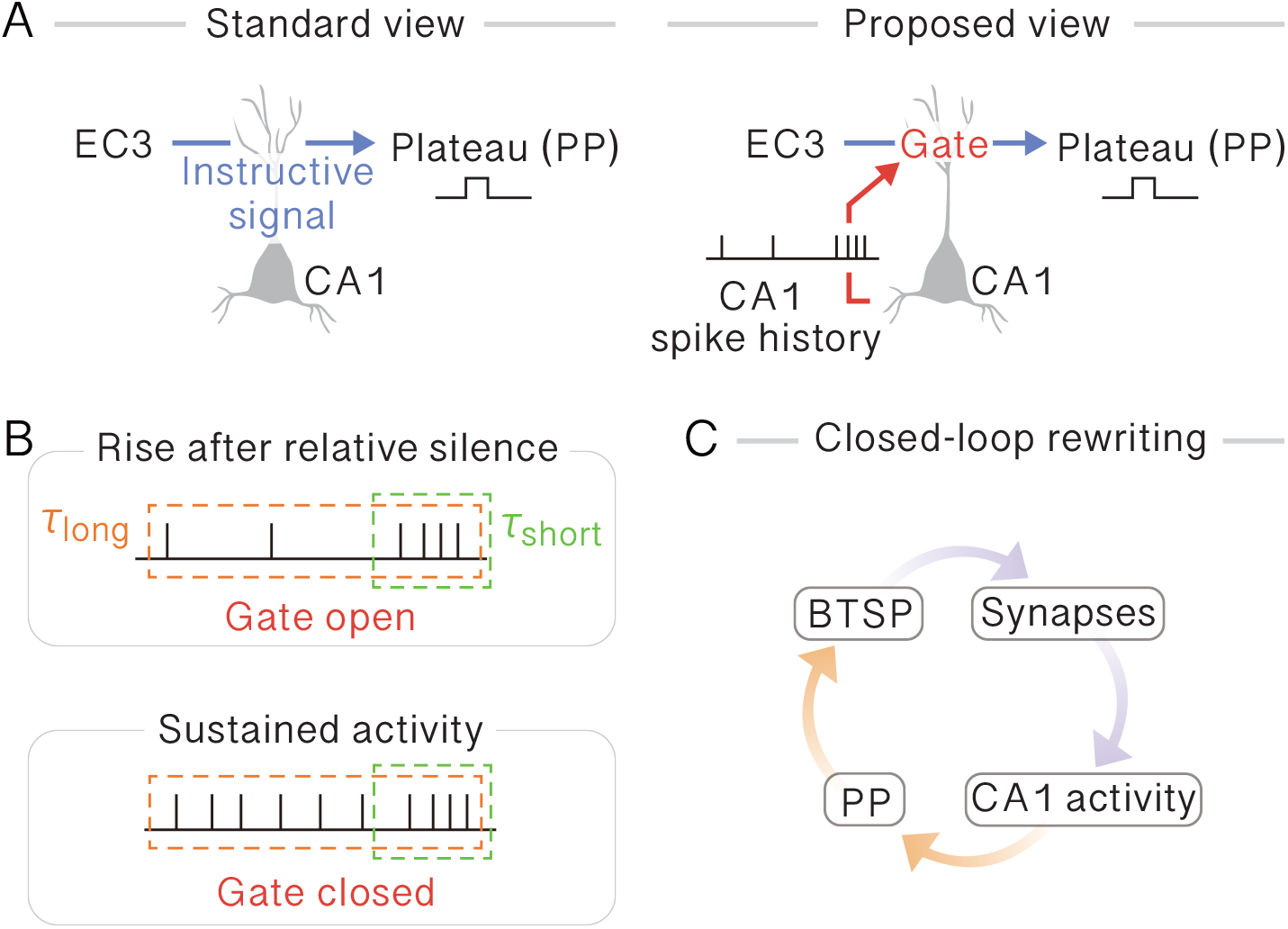
Spike-history-dependent plateau gating enables closed-loop representational rewriting. **A** Schematic showing two levels of control over PP initiation. EC3 provides a target-like drive of PPs (left), while recent CA1 spike history controls whether an individual neuron is available for PP initiation (right). **B** Spike-history-dependent gating distinguishes a rise in firing after relative silence from sustained activity. When a burst of spikes follows relative silence, activity in the recent window is high while activity over the preceding longer window remains low, opening the gate. During sustained activity, recent spiking can be similarly high, but elevated activity in the longer window keeps the gate closed. *τ*_short_ and *τ*_long_ indicate the recent and longer spike-history windows, respectively. **C** Closed-loop interaction between synaptic state, CA1 activity, PPs, and BTSP.

To address this question, we propose that recent somatic spiking history provides a means for cell-specific control of plasticity events (Figure 1A, right). In this framework, EC3 biases plateau-potential (PP) generation, while recent spike history gates when the postsynaptic CA1 neuron is in a state that allows for plasticity events. The gate distinguishes a rise in firing after relative silence from sustained activity (Figure 1B), consistent with the recently observed spike-rate accelerometer mechanism [21, 22]. Specifically, the gate opens for a short period when the number of somatic spikes in a brief recent window exceeds a threshold while activity in a preceding longer window remains low. The same level of recent spiking does not open the gate when it was embedded in sustained activity. EC3 therefore modulates PP probability as a permissive signal only when the spike-history gate is open, linking broad modulation to cell-specific plasticity events.

This history-dependent control creates a cyclic interaction between representations and plasticity (Figure 1C). The current synaptic state shapes CA1 activity, and the resulting activity history biases where a subsequent PP can occur. Each PP then triggers behavioral time-scale synaptic plasticity (BTSP), which modifies the synaptic state and thereby changes the activity pattern that will constrain future PPs. Thus, the current representation helps determine when it is rewritten next, closing a feedback loop between neural activity and synaptic plasticity.

### 2.2 CA1 activity rather than EC3 input can specify when plasticity occurs

To test whether this principle allows for and is consistent with spatial representation in hippocampal pyramidal cells, we implemented the spike-history gate in a two-compartment spiking neuron model of CA1 (Methods). PP initiation depended jointly on the spike-history gate and EC3 input. Each PP triggered BTSP at CA3-to-CA1 synapses and was followed by a refractory period during which further PPs were suppressed.

We then asked whether an existing place field biases subsequent plasticity as an agent repeatedly traversed a 300-cm circular track at constant velocity (Figure 2A, left). Model CA1 neurons received spatially tuned CA3 input onto proximal dendrites [5, 18] and spatially unstructured, stationary EC3 input following Grienberger and Magee [13] onto distal apical dendrites (Figure 2A, right). Plasticity at CA3-to-CA1 synapses followed a weight- and timing dependent bidirectional BTSP rule, allowing for both potentiation and depression of synapses (Figure S1). CA1 model neurons also interacted through lateral inhibition at the soma to avoid spatial clustering of place fields [23, 24]. Place fields emerged within approximately five to ten laps and shifted gradually over continued trials (Figure 2B). A transient novelty modulation accelerated initial field formation but was not required for (Figure S2), and the model reproduced the approximately linear dependence of place-field size on running velocity [4] (Figure S3).

**Figure 2:**
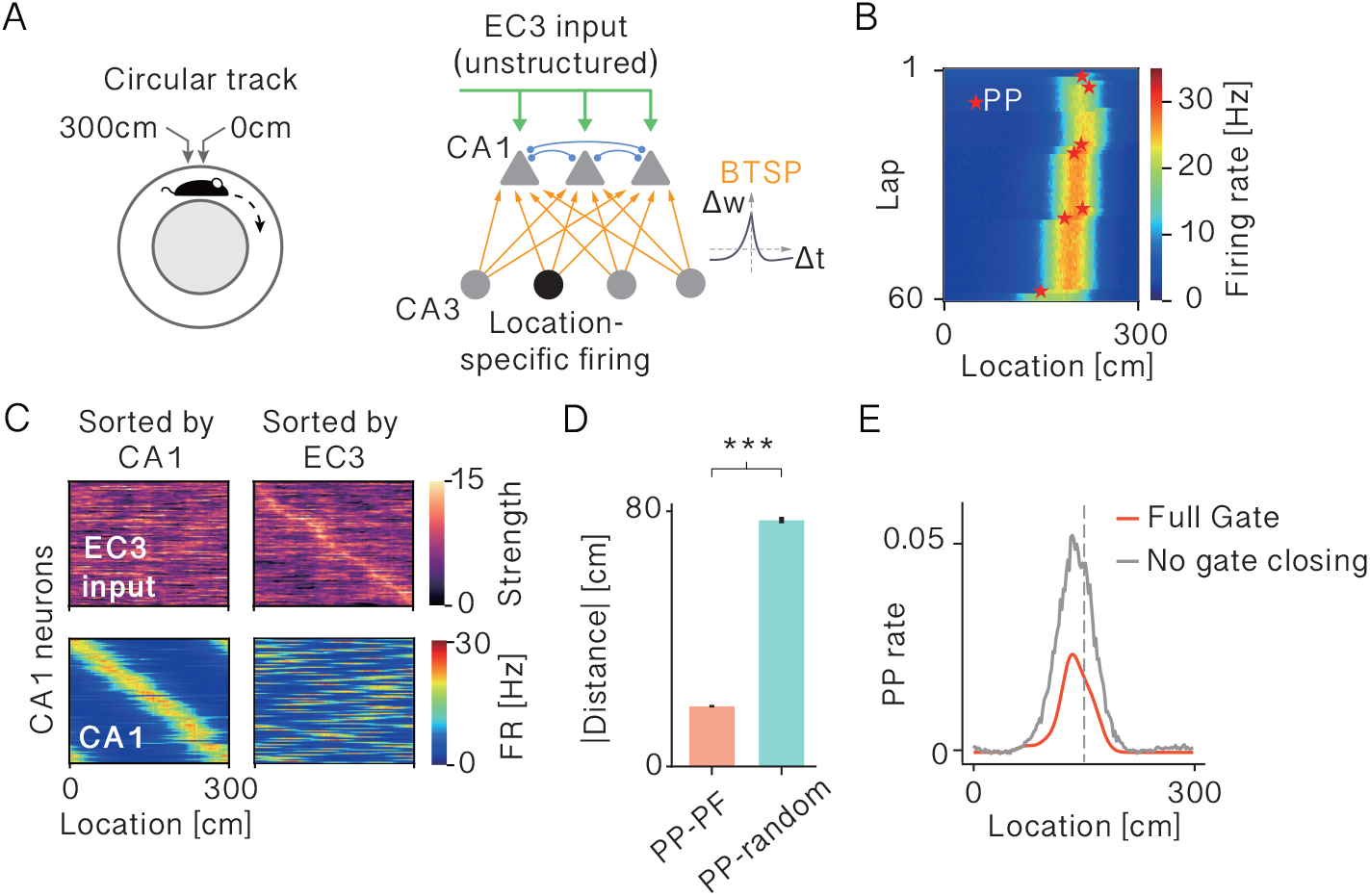
Existing place fields bias where subsequent plateau potentials occur. **A** Simulation setup. An agent repeatedly ran along a 300-cm circular track (left). CA1 model neurons received spatially tuned CA3 input and spatially unstructured EC3 input (right). Plateau potentials triggered BTSP at CA3-to-CA1 synapses. **B** Place-field evolution of an example CA1 model neuron across 60 laps. Red stars indicate plateau potentials (PPs). **C** Example EC3 input to CA1 neurons (top) and CA1 population firing rates (bottom). Neurons are shown either sorted according to their CA1 place-field locations (left) or according to their EC3 inputs (right). **D** Average median distance between the location of each PP and the place-field center of the same neuron on the preceding lap, compared with the distance between the PP and a random reference location. **E** Spatially resolved PP rate of model neurons with and without PP gate closing mechanism. The place field was induced at the center of the track (vertical dashed line). In **D**, P values were obtained from one-sided Welch’s t-tests across 5 independent simulations (*∗ ∗ ∗p <* 0.001).

We next visualized the relationship between the emerging CA1 spatial representation and the corresponding EC3 input. Sorting neurons by their CA1 place-field locations revealed a clear progression across the track while the corresponding EC3 inputs remained unstructured (Figure 2C, left). Conversely, sorting the same neurons according to their EC3 inputs did not reveal a corresponding spatial order in CA1 activity (Figure 2C, right). Thus, the CA1 spatial sequence was not accompanied by a matching fine-grained spatial organization in the EC3 input.

Given that the CA1 spatial organization was not directly inherited from EC3 input, we then wondered whether the current CA3-driven place-field representation could instead bias where the history-dependent gate opened. We quantified the distance between each PP and the place-field center of the same neuron on the preceding lap and compared it with the distance from the PP to a random reference location drawn from a uniform distribution. Subsequent PPs occurred closer to the previous place-field center than to the random reference, showing that the current representation spatially biased the next plasticity-inducing event (Figure 2D). To test the contribution of the preceding low-activity requirement, we removed this component while retaining the dependence on elevated recent firing. This manipulation drastically increased PP rate without substantially shifting the spatial peak of PP occurrence relative to the expressed place field (Figure 2E). Thus, elevated recent firing is sufficient to preserve the spatial preference of PP initiation, whereas the preceding activity history determines whether that firing represents a new opportunity for plasticity or sustained activity that should be suppressed. By coupling PP initiation and current place field expression via its recent spiking activity, a neuron may thus avoid repeated and putatively costly plasticity events at locations that are already strongly expressed.

Together, these results show that, in the proposed framework, EC3 input need not specify the fine-grained spatial organization of CA1 activity. Instead, the current place-field representation biases where the spike-history gate opens and constrains where subsequent PPs occur, allowing repeated local updates to produce gradual representational drift without disrupting the population code (Figure S4, Figure S5).

### 2.3 BTSP Closes the Feedback Loop Between Place-Field Activity and Synaptic State

We next asked how PPs and the subsequent plasticity update the synaptic state leading to place field drift. We focused on a model neuron whose place field drifted smoothly over repeated laps (Figure 3A), allowing us to follow the underlying CA3-to-CA1 synaptic changes as the represented location moved. By the final lap, the CA3-to-CA1 weight profile was concentrated around the location of the place field expressed at that time (Figure 3B), indicating a close correspondence between the synaptic weight structure and the current place-field representation.

**Figure 3:**
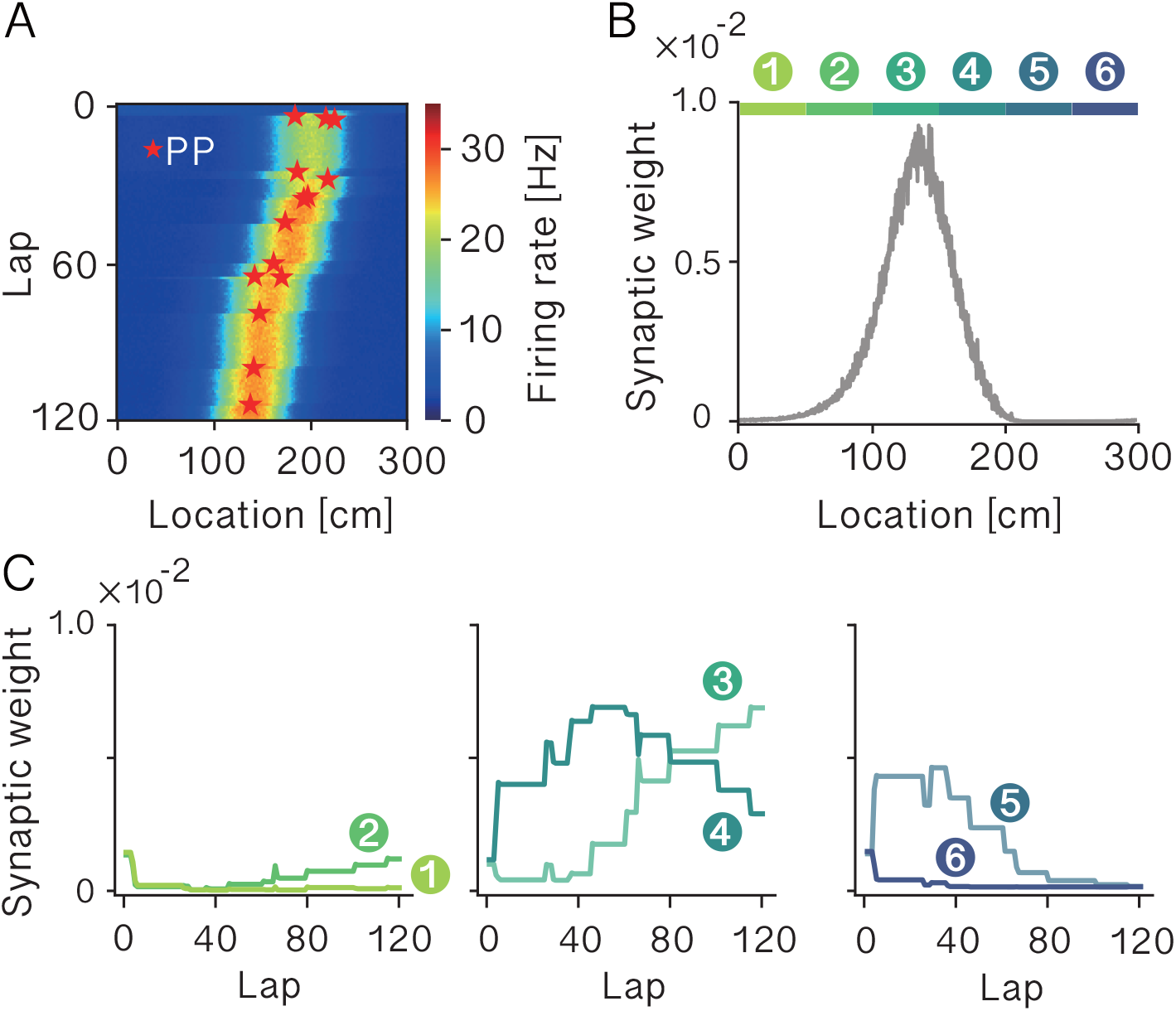
Synaptic weight dynamics underlying place-field drift. **A** Place-field evolution of an example CA1 model neuron across 120 laps. Red stars indicate plateau potentials (PPs). **B** Final CA3-to-CA1 synaptic weights after the penultimate lap. CA3 inputs are ordered according to their place-field locations along the track. Colored bars indicate the six spatial regions used for the analysis in C. **C** Evolution of the average synaptic weight within each of the six regions indicated in B. Different groups of CA3 inputs underwent successive potentiation and depression as the CA1 place field drifted across the track.

To determine how this synaptic profile emerged, we divided the ordered CA3 inputs into six spatial regions and tracked the mean weight within each region across laps (Figure 3B, C). As the drifting field approached a region, inputs representing that region were consistently potentiated; after the field moved beyond it, the same inputs weakened again. Different groups of CA3 inputs therefore passed through successive phases of potentiation and depression as the place field moved along the track. Synapses supporting upcoming field locations gained strength, whereas synapses associated with locations left behind subsequently weakened. The synaptic support for the place field thus translated across the CA3 input population together with the drifting representation.

Altogether, these dynamics close the proposed feedback loop between representation and plasticity. The current CA3-to-CA1 configuration shapes the place-field activity that biases where the spike-history gate can open, while each resulting PP modifies that same synaptic configuration through BTSP. The synaptic state is therefore continuously updated as the representation drifts [5, 18].

### 2.4 Broad EC3 Reward Modulation Biases Cell-Specific Plasticity

Given a history-dependent gating of PPs, what is the effect of broad EC3 modulation on plasticity induction? Reward provides a natural test case for this scenario. CA1 place fields are overrepresented near reward sites [25, 26], and EC3 activity is known to contribute to this organization. Indeed, inhibiting EC3 axons near reward prevents place-field overrepresentation, while EC3 activity and axon-to-axon correlation increase near reward [13]. We therefore asked whether broad reward-related EC3 modulation could bias CA1 representations without specifying the target location of individual cells.

To test this, we simulated an agent running at constant velocity on a 300-cm circular track with a central reward site (Figure 4A). EC3 input was sampled from a non-stationary process with elevated activity around reward, modeled after the changes reported by Grienberger and Magee [13]. Running behavior was held fixed (constant velocity), allowing us to isolate the effect of EC3 modulation itself. We found that broad reward-related EC3 modulation was sufficient to bias the CA1 spatial representation toward the reward location. In the reward condition, CA1 place fields became concentrated around the reward location, coinciding with the broad increase in EC3 input received by the corresponding CA1 neurons (Figure 4B). PP rate was elevated around the same reward location (Figure 4C). Broad EC3 modulation therefore increased opportunities for BTSP near reward, while the current CA3-driven state of each neuron continued to constrain the precise location at which plasticity occurred. Consistent with this redistribution, population-averaged CA1 activity was also elevated around the reward location relative to the no-reward condition (Figure 4D).

**Figure 4:**
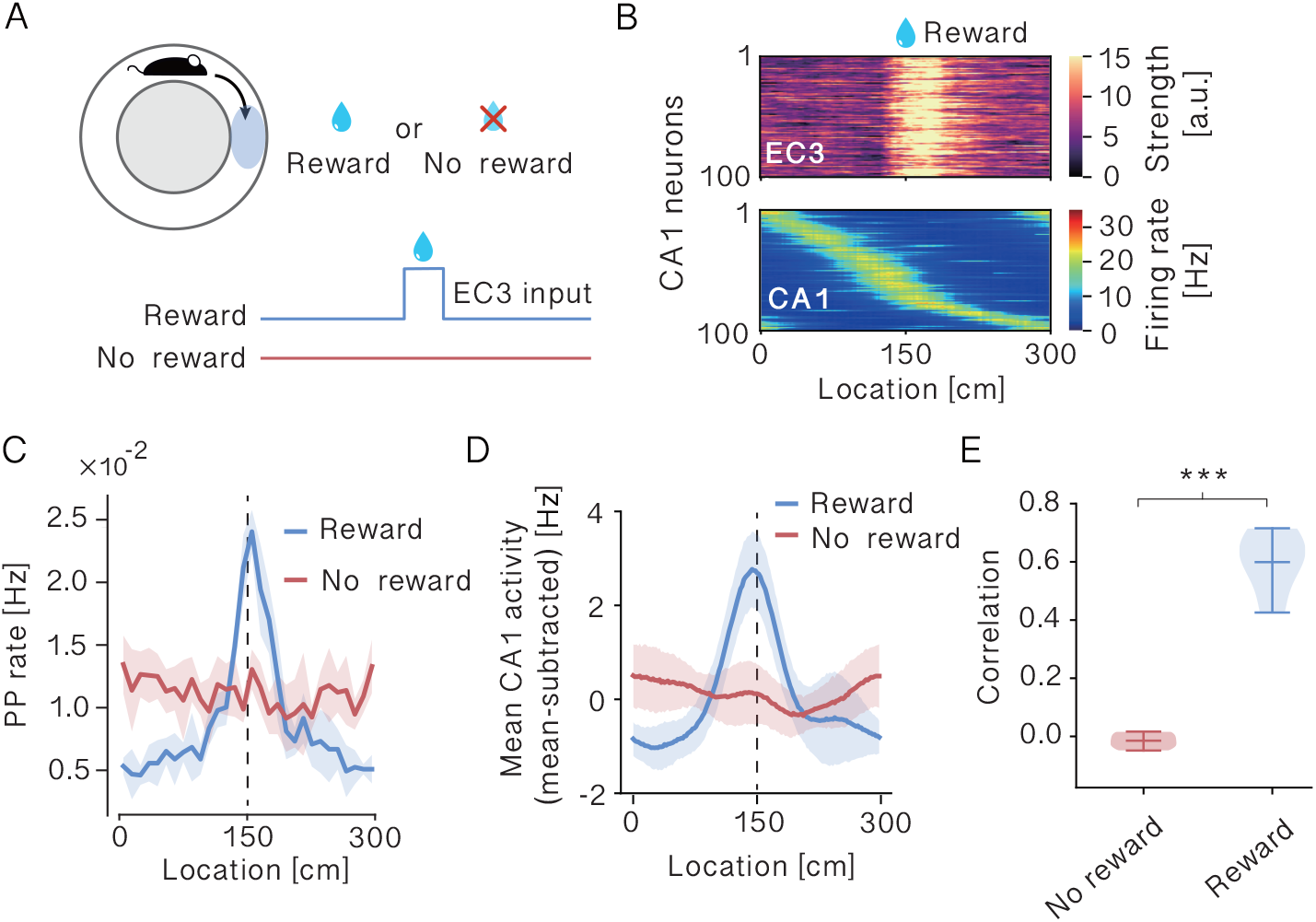
Reward-modulated EC3 input biases plateau-potential initiation and CA1 representations. **A** Schematic of the reward manipulation. The agent ran at constant velocity along a 300 cm circular track. In the reward condition, EC3 input was transiently increased around the reward location at 150 cm for 1.5 s, whereas EC3 input remained spatially stationary in the no-reward condition. **B** Example EC3 input to CA1 model neurons (top) and CA1 population firing rates (bottom) in the reward condition. CA1 neurons are sorted by the location of their peak firing, and the same neuron order is used to display the corresponding EC3 inputs. **C** Spatially resolved plateau-potential (PP) rate in reward and no-reward conditions. The dashed line indicates the reward location. **D** Spatial profile of population-averaged CA1 activity in reward and no-reward conditions. In C and D, lines and shaded regions indicate the mean ± SD across five random seeds. **E** Pearson correlation between summed CA1 population activity and summed EC3 input across the track in no-reward and reward conditions. In E, P values were obtained from one-sided Welch’s t-tests across 5 independent simulations (*∗ ∗ ∗p <* 0.001).

Reward modulation also reproduced population-level coupling between CA1 activity and EC3 input. Summed CA1 activity and EC3 input were essentially uncorrelated in no-reward runs but strongly positively correlated in reward runs (Figure 4E). The correlation between mean CA1 population activity and mean EC3 input likewise rapidly increased and remained stable across laps (Figure S6), consistent with experimental data (Grienberger and Magee [13], their Figure 6a).

Together, these results show that broad EC3 modulation can bias PP initiation and CA1 representations toward reward at the population level. The precise location of plasticity in each neuron, however, remains constrained by its current CA3-driven state.

### 2.5 Previous Representations Remain Recoverable After Remapping

CA1 place-field representations can reorganize rapidly between environments yet remain partially recoverable upon return to a familiar environment [27, 28] (but see [29]). We therefore asked how substantial synaptic plasticity could occur in one environment without necessarily erasing the representation expressed in another.

We simulated a 300-cm circular track in two environments (Figure 5A). To model partially overlapping CA3 representations across environments, we used two CA3 populations with 50% overlap [28, 30]. For each environment, the place-field centers are drawn independently. The agent first experienced environment 1 for 20 laps, then spent a variable number of laps in environment 2, and finally returned to environment 1. When the agent re-entered environment 1, its previous CA1 representation was rapidly re-expressed, whereas the representation expressed in environment 2 was decorrelated from it (Figure 5B). Thus, intervening plasticity in environment 2 did not simply overwrite the activity pattern associated with environment 1. Full cross-sorting comparisons of the population activity maps are shown in Figure S7.

**Figure 5:**
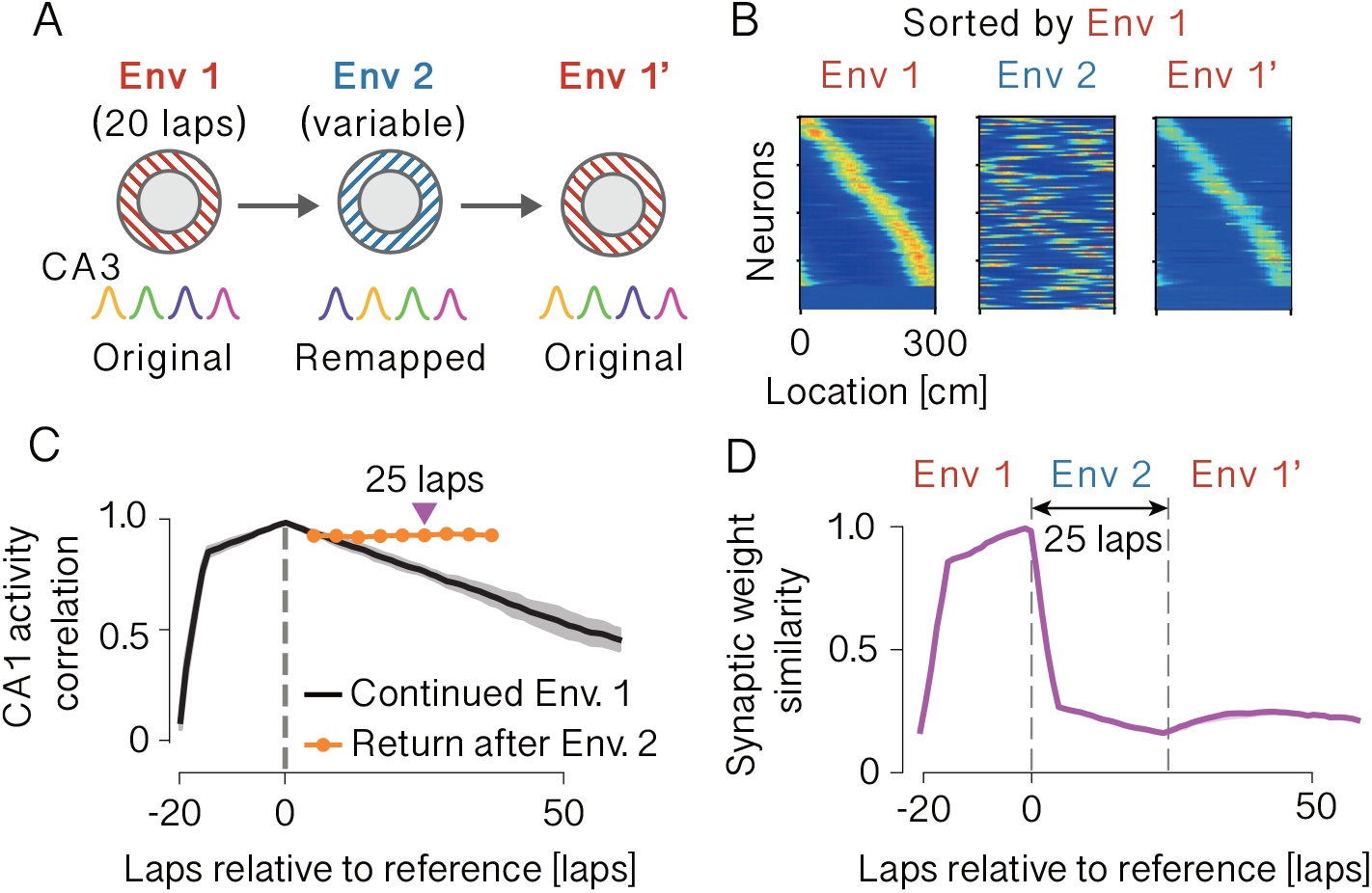
Previous representations remain recoverable despite synaptic changes during remapping. **A** Simulation protocol. The agent first experienced environment 1 (Env 1) for 20 laps, followed by a variable number of laps in environment 2 (Env 2), and then returned to Env 1 (Env 1^*′*^). The same CA3 neurons were used across environments, but their place-field tuning was remapped in Env 2 and restored in Env 1^*′*^. **B** Example CA1 population activity in Env 1, Env 2, and after return to Env 1^*′*^. Neurons are sorted according to their place-field locations in Env 1. The Env 1 representation was disrupted in Env 2 but re-emerged upon return to Env 1^*′*^. **C** CA1 population activity correlation relative to the final lap of the initial Env 1 exposure. Continued exposure to Env 1 produced progressive decorrelation, whereas correlations measured upon return to Env 1 after intervening Env 2 exposure remained comparatively high. The triangle marks the 24-lap Env 2 condition shown in D. **D** CA3-to-CA1 synaptic weight similarity for the representative 25-lap Env 2 condition, measured relative to the same reference lap as in C. Synaptic similarity decreased during Env 2 exposure despite the relative preservation of the Env 1 representation shown in C, indicating that substantial synaptic change during Env 2 need not erase the previous representation.

**Figure 6:**
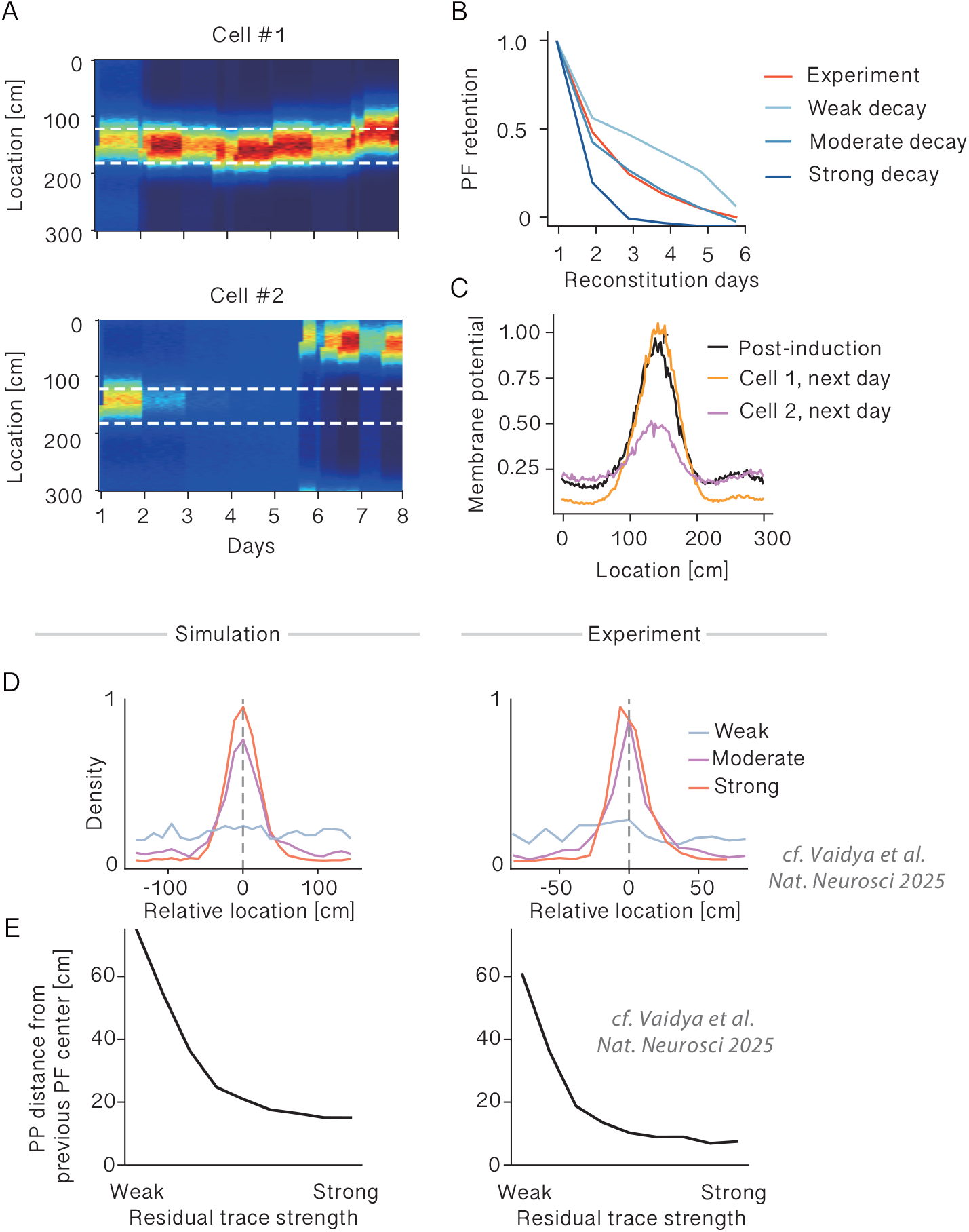
Residual synaptic structure biases place-field reconstitution across days. **A** Place-field activity of two example CA1 model neurons across simulated days. Cell #1 repeatedly formed a place field near its previous location, whereas Cell #2 eventually formed a place field at a different location. Dashed lines indicate the spatial range around the initially induced place field. **B** Place-field retention across days in the experimental data and in simulations with different strengths of between-day synaptic decay. **C** Membrane-potential profiles immediately after place-field induction and on the following day for the two example neurons in A. **D** Distribution of subsequent plateau-potential (PP) locations relative to the previous place-field center in the simulation (left) and experimental data (right). For the simulation, the weak, moderate, and strong residual-trace conditions correspond respectively to the strong, moderate, and weak synaptic-decay conditions shown in B. Stronger residual traces produced a sharper concentration of subsequent PPs around the previous place-field location. Experimental data showed the same relationship when grouped according to the strength of pre-plateau residual activity around the previous place field. **E** Distance of subsequent PPs from the previous place-field center as a function of residual trace strength in the simulation (left) and experimental data (right). In both cases, stronger residual traces were associated with smaller PP distances from the previous place-field location.

We next quantified how the environment 1 representation changed with experience. Taking the final lap of the initial environment 1 exposure as a reference, continued exposure to environment 1 produced progressive decorrelation (Figure 5C, black graph). In contrast, correlations measured upon return to environment 1 after intervening exposure to environment 2 remained comparatively high, even as the duration of the intervening exposure increased (Figure 5C, orange graph). The complete correlation dynamics across the initial Env 1 exposure, intervening Env 2 exposure, and return to Env 1 are shown in Figure S8. After reentering environment 1 the spatial representation continued to drift at a similar rate as without environment change (Figure S9). These results are consistent with the recently reported experience dependence of the spatial representation [31, 32] that show that drift was controlled primarily by time of experience in the corresponding environment rather than by elapsed time.

Synaptic weights nevertheless continued to change substantially during environment 2. For the representative 24-lap environment 2 condition highlighted in Figure 5C, CA3-to-CA1 weight similarity rapidly decreased during the intervening exposure even though the environment 1 activity pattern remained comparatively well preserved upon return (Figure 5D). Upon entering environment 1 weight similarity initially increased again, suggesting a partial recovery of rewritten weights. The dissociation between substantial synaptic change, modest loss of the environment 1 representation, and weights recovery shows that the functional impact of plasticity depends on the input pattern of CA3 activity. This is consistent with the idea that synaptic changes that are weakly aligned with the CA3 input pattern used in environment 1 have only limited effects on the corresponding representation.

In summary, substantial synaptic changes in one environment did not necessarily erase the representation expressed in another. The same evolving synaptic weights could therefore support partial recovery of distinct representations depending on the current CA3 input pattern.

### 2.6 Surviving Synaptic Structure Biases Place-Field Reinstatement Across Days

We finally asked whether the same feedback principle could account for the recently observed cross-day reconstitution of place fields. Vaidya et al. [33] showed that CA1 place fields can reform at similar locations across days despite substantial loss of learned synaptic strength and suggested that residual place-field activity may bias the probability and location of subsequent plateau potentials. This raises a mechanistic question: what allows a previous place-field location to influence where the next plateau potential occurs?

We modeled this process in individual CA1 neurons without lateral inhibition and induced an initial PP at the track midpoint. Between simulated days, CA3-to-CA1 weights were partially relaxed toward their initial values to mimic incomplete retention of learned synaptic changes across days. To isolate cross-day re-formation from repeated within-day plasticity, we reduced the probability of additional PPs after the first PP on each simulated day.

Individual neurons showed different cross-day outcomes depending on how much location-specific structure remained. Cell #1 repeatedly formed a place field near its previous location, whereas Cell #2 eventually formed a field at a different location (Figure 6A). We next varied the strength of between-day synaptic decay and compared place-field retention (assessed by the difference of place field centers across days) with the experimental data of Vaidya et al. The rate of cross-day retention depended strongly on decay strength, with an intermediate decay regime providing the closest match to the experimentally observed decline (Figure 6B).

We next examined the residual membrane-potential profiles of the two example neurons shown in Fig. 6A. Although between-day decay weakened the learned CA3-to-CA1 weights, it did not erase them completely. In Cell #1, which repeatedly formed a place field near its previous location, the remaining synaptic structure produced a clear residual depolarization around that location on the following day. This residual depolarization was weaker in Cell #2, which subsequently formed a place field elsewhere (Figure 6C). Thus, even after the previous place field had weakened, its underlying synaptic structure could leave a spatially localized trace in membrane potential.

The strength of the residual trace also predicted where subsequent PPs occurred. In the model, stronger residual traces produced a sharper concentration of PPs around the previous place-field location, and PP distance from the previous place-field center decreased with increasing residual trace strength (Figure 6D, E, left). Reanalysis of the Vaidya et al. [33] data revealed the same relationship: stronger pre-plateau residual activity was associated with subsequent PPs occurring closer to the previous place-field location (Figure 6D, E, right). Thus, the model and experimental data showed the same spatial dependence between the residual trace and subsequent PP location, a relationship not directly examined in the original study.

In summary, partial synaptic decay converts continuous place-field drift into cross-day reformation while preserving enough location-specific structure to bias where that reformation occurs. Surviving synaptic structure leaves a residual activity trace that biases subsequent PPs toward the previous place-field location, allowing BTSP to reconstitute a similar representation across days.

## 3 Discussion

The central problem investigated in this study is how an individual neuron can remain sensitive to weak but meaningful activity that should reinforce an existing representation, while limiting unnecessary plasticity once that representation is strongly expressed. This creates a sensitivity-stability dilemma for the timing of plateau potentials in BTSP. The spike-rate accelerometer identified by Park et al. [21] and Lee et al. [22] provides a temporal asymmetry required to address the problem. A burst after relative silence strongly boosts back-propagating activity and distal dendritic depolarization, whereas sustained activity does not and even prevents such a boost. We therefore interpret recent spike history as a cell-specific gate for plateau initiation. Madar et al. [18] showed that probabilistic BTSP induction triggered by neural spiking can account for gradual place-field drift. However, a firing-dependent rule alone does not specify how a neuron distinguishes newly rising activity from sustained firing at an already expressed field. Our model provides a candidate cellular mechanism by comparing recent spiking with the preceding activity history. We predict that a sharp rise in firing after relative silence can promote plateau initiation even when a place field is already present, whereas sustained firing of the same magnitude should suppress it.

Once a PP occurs, BTSP changes CA3-to-CA1 weights and therefore the activity pattern that constrains future PP locations. Repetition of this feedback loop produces gradual drift while preserving spatial coverage. Its functional effect depends on the input pattern through which the modified weights are read out, allowing representations to remain recoverable after plasticity in another environment. Across days, partial retention of the same synaptic state leaves a location-specific bias that can guide a new PP and restart BTSP. Drift and cross-day stability are therefore different regimes of the same cyclic process, set by how strongly the preceding synaptic structure is retained and expressed.

The model also bears on the proposal that BTSP implements a form of target learning in CA1, in which EC3 conveys a target signal that CA3-to-CA1 plasticity learns to approximate [13, 15]. At the population level, such a signal can specify where learning is favored, but it leaves a cell-specific credit-assignment problem: an individual CA1 neuron still requires a mechanism that determines whether its current activity should trigger plasticity. Individual EC3 axons are broadly tuned and variable across trials, and EC3 modulation predicts where plasticity is enhanced without determining individual place-field locations. Likewise, oriens-lacunosum moleculare (OLM) interneuron activity rises over learning and suppresses further PPs broadly across the population [34, 35]. These observations are consistent with population-level instructive control while leaving the assignment of plasticity to individual cells unresolved. Our framework supplies this missing cell-specific factor. Recent somatic history determines when the PP gate can open, the current CA3-driven state constrains where it opens, and EC3 controls the probability that a PP is generated once the gate is permissive. Thus, EC3 acts as a permissive target signal that biases where and how strongly learning is favored, while each cell’s recent activity history determines whether its own state is eligible for rewriting.

The origin of representational drift has often been considered in terms of ongoing synaptic turnover, parameter fluctuations, or diffusion through a space of functionally equivalent representations [36, 37, 38]. Similar to Madar et al. [18], in our model, drift instead arises from structured plasticity that is conditioned on the representation currently being expressed. Because subsequent PPs are biased toward the current place field, each plasticity event locally rewrites the existing representation rather than displacing it through an unconstrained random walk like a L’evy Flight. This provides a direct explanation for why spatial drift depends on active experience in the corresponding environment, as observed experimentally [31, 32].

Several open questions remain. The history-dependent gate abstracts the accelerometer mechanism and does not reproduce the measured quantitative dependence of dendritic excitability or plateau probability on preceding spikes. EC3 is modeled phenomenologically, and its real activity is more heterogeneous than the spatially unstructured baseline and broad reward modulation used here. What is the role of the inhibitory circuit in controlling the induction of plasticity events (see [24, 35, 34, 39, 40])? We also omit partial remapping structure, variable running speeds, and theta-scale timing between CA3 and EC3 inputs, which could interact with the proposed gate [12]. Finally, the Vaidya et al. [33] reanalysis is correlational: residual activity and subsequent plateau localization could both reflect recurring spatial input rather than a causal effect of the residual trace itself. These simplifications define direct tests for more physiological implementations and experiments.

Our framework provides several experimentally testable predictions. Whole-cell voltage imaging or intracellular recordings should reveal a location-specific subthreshold depolarization at the previous field location before place-field reconstitution, whose amplitude predicts how strongly the next plateau potential is spatially biased toward that location. Imposed activity should gate plasticity according to its history: onset-like transients after relative silence should permit plateau initiation, whereas sustained activation of the same magnitude should suppress it. Similar to the findings reported by Noguchi et al. [41], future research should investigate differential roles for proximal and apical dendrites as drivers of plasticity events in CA1 during reinstantiation or remapping. Our results suggest inputs to proximal dendrites as locus for reinstantiating and reinforcing plasticity events.

More generally, the proposed history dependence provides a way to separate the expression of an existing representation from its subsequent update. This view is also consistent with previous work assigning computational roles to somatodendritic interactions [42]. By making plateau initiation sensitive to a rise from relative silence rather than sustained firing, recent somatic history determines when plasticity becomes possible, while the current CA3-driven representation constrains where the gate can open and EC3 biases whether a PP is generated. This feedback links future plasticity to the state already stored in CA3-to-CA1 synapses and unifies within-environment drift, context-dependent recoverability, reward-associated overrepresentation, and cross-day reconstitution. Given that BTSP-like plasticity extends beyond the hippocampus, the same principle may allow fast event-triggered plasticity to remain coupled to changes in neural state without being continuously re-engaged by the state itself.

## 4 Data Availability

The neural recording data from Vaidya et al. [33] are available on Figshare (https://doi.org/10.6084/m9.figshare.29052116.v1). The synthetic data used in this study can be reproduced using the source code.

## 5 Code availability

The complete source code used for the numerical simulations and data analyses will be made publicly available upon publication.

## 6 Author Contributions

ACK and TA conceived the study and wrote the paper. ACK performed the simulations and data analyses.

## 8 Acknowledgments

This work was supported by RIKEN Center for Brain Science (TA) and RIKEN Cluster for Pioneering Research (TA). We are grateful to Yoshihito Saito for fruitful discussions.

## 9 Methods

### 9.1 Spike-History-Dependent Gating in a Two-Compartment Neuron Model

CA1 model neurons have a somatic compartment targeted by CA3 inputs and a dendritic compartment targeted by EC3 input. We integrated incoming CA3 model neuron spikes linearly according to

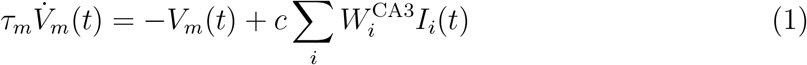

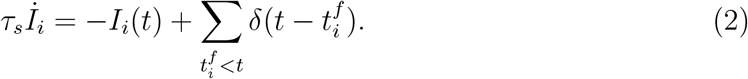

Here, *τ*_*m*_ is the membrane time constant, *V*_*m*_ is the somatic membrane potential, *c* a scaling factor (depending only on the time constants), 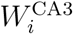 denotes synaptic weights of CA3 connections, *τ*_*s*_ is the synaptic time constant, *I*_*i*_ the synaptic current evoked by a single CA3 model neuron, and 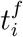 are the spikes emitted by CA3 model neurons. We integrate the membrane potential using an exponential integrator [43]. Somatic spikes are sampled using the instantanoues firing rate [44]

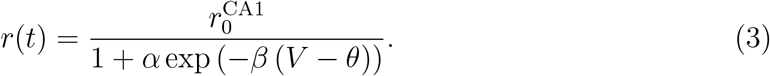

The apical compartments receive modulatory input from EC3 model neurons of the form

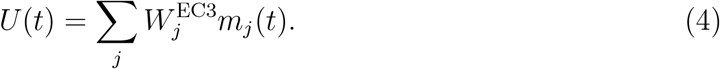

Here, *W* ^EC3^ are the synaptic weights of EC3 connections, and *m*_*j*_(*t*) the activity of EC3 model neurons. Each CA1 model neuron has an additional binary gating variable *G*(*t*) initially set to 0. *G*(*t*) depends on its own spiking history through the intervals (*t − τ*_short_, *t*] and (*t − τ*_long_, *t*]. If the total number of spikes in the first interval exceeds *n*_low_ and the total number of spikes in the second interval is less than *n*_high_, then the gating variable is set to 1 for *τ*_window_. Otherwise, *G*(*t*) = 0.

Plateau potentials are sampled from an inhomogeneous Poisson process with instantaneous rate

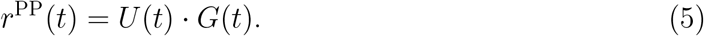

Once a PP is sampled, no new PP can be generated for *τ*_PP*−*refac_. Single neuron parameters are displayed in Table 1. The scaling constant *c* is defined as

**Table 1:** Parameters specifying CA1 model neuron dynamics.

| Single Neuron Parameters |  |  |
| --- | --- | --- |
| Parameter | Value | Meaning |
| $\tau_m$ | 20 ms | Membrane time constant |
| $\tau_s$ | 5 ms | Synaptic time constant |
| $\alpha$ | 0.5 | Rate-sigmoid scaling |
| $\beta$ | 3 | Rate-sigmoid gain |
| $\theta$ | 1 | Soft spiking threshold |
| $r_0^{\text{CA1}}$ | 30 Hz | Maximal firing rate |
| $\tau_{\text{short}}$ | 50 ms | PP trigger window |
| $\tau_{\text{long}}$ | 200 ms | PP closing window |
| $n_{\text{low}}$ | 3 | Spike count trigger |
| $n_{\text{high}}$ | 6 | Spike count trigger |
| $\tau_{\text{window}}$ | 100 ms | PP sampling window |
| $\tau_{\text{PP-refac}}$ | 1000 ms | PP refractory |

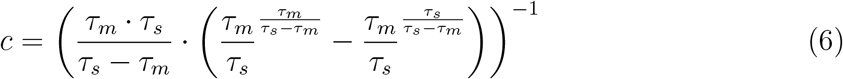

### 9.2 Behavioral Time Scale Plasticity

We updated the synaptic weights of the CA3 model neurons with a BTSP model. The updates depended on both, the timing of PPs, and the presynaptic activity. For this, we computed four plasticity traces. First, for each synapse the trace depended on the activity of the presynaptic neuron *i* according to:

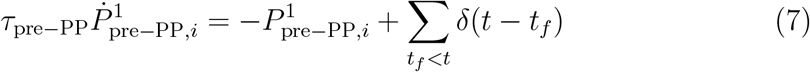

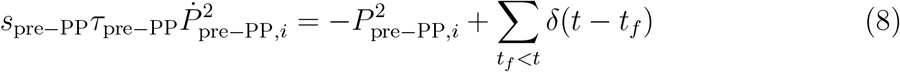

Similarly, for each CA1 model neuron, we computed the traces depending on the PP timing *t*_PP_ as

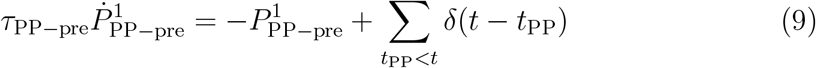

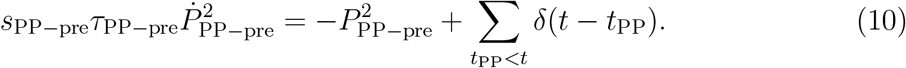

Upon a presynaptic spike of neuron *i* at time *t*_*f*_, the weight was updated according to

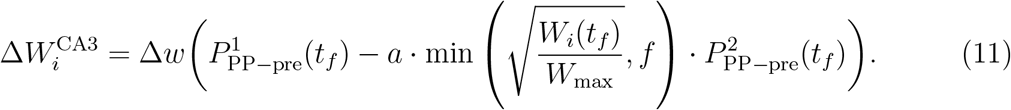

When a PP occurs in the postsynaptic neuron at time *t*_PP_, the weight of presynaptic neuron *i* was updated according to

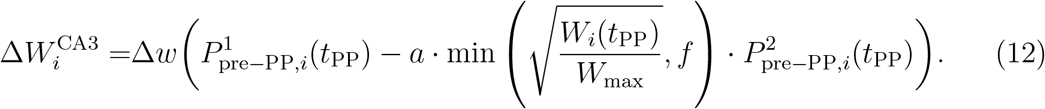

The plasticity kernel is thus a difference of exponentials, where the contribution of the subtracted exponential is controlled by the size of the weight before the potentiation or depression. The weights are limited to not exceed *W*_max_ or fall below 0. Plasticity parameters are displayed in Table 2.

**Table 2:**
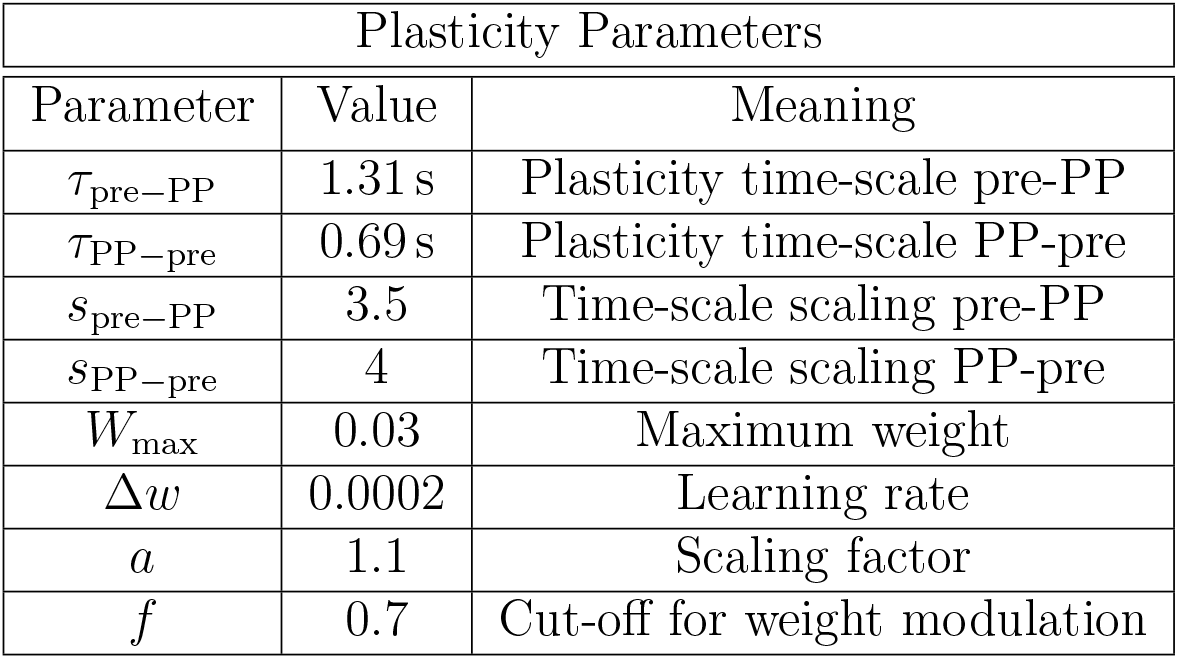
Parameters specifying weight updates.

### 9.3 Network model

The simulated networks consist of *N*_CA1_ CA1 model neurons that receive inputs to their somatic compartments from *N*_CA3_ CA3 model neurons and apical input from *N*_EC3_ EC3 model neurons. The weights *W* ^CA3^ are independent and identically distributed (i.i.d), and drawn from a clipped normal distribution with mean *µ*_CA3_ and standard deviation *σ*_CA3_, preventing negative weights. *W* ^EC3^ are i.i.d. sampled from a Bernoulli distribution with connection probability *p*^EC3^ and with positive weight *w*^EC3^. The CA1 model neurons are recurrently connected with i.i.d. weights sampled from a Bernoulli distribution with connection probability *p*^rec^ and with weight *−w*^rec^ (excluding self-connections). In our model, recurrent interactions have an inhibitory effect, representing the di-synaptic inhibition of recurrent connections in CA1. Recurrent spikes are integrated linearly akin to spikes originating from CA3 model neurons. At the beginning of each numerical experiment the total sum of CA3 weights to a single CA1 model neuron is normalized to 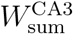 exactly once.

CA3 model neurons sample spikes from an inhomogeneous Poisson process. Each neuron has a Gaussian shaped place field, and their firing rate depends on the location *x* of the agent on the track:

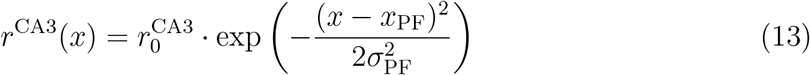

Here, *x*_PF_ determines the center of a place field and *σ*_PF_ its width. The place fields of the CA3 model neurons cover the circular track evenly with equidistant place field centers. In the case where the agent experiences multiple environments there are two sets of *N*_CA3_ model neurons with a 50% overlap. For each set, the place field centers are drawn independently.

EC3 neurons are modeled as suggested by Grienberger and Magee. In short, EC3 activity is modeled by binary neurons. Their activity *m*(*t*) switches between ON and OFF states according to a Markov-chain model with transition matrix

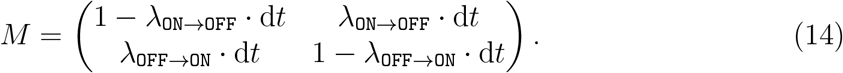

The state is updated at each time step.

All network parameters are displayed in Table 3. Parameters for CA3 input can be found in Table 4, and EC3 input parameters can be found in Table 5.

**Table 3:** Parameters specifying network.

| Network Parameters |  |  |
| --- | --- | --- |
| Parameter | Value | Meaning |
| $N_{\text{CA1}}$ | 100 | Num. of CA1 model neurons |
| $N_{\text{CA3}}$ | 1500 | Num. of CA3 model neurons |
| $N_{\text{EC3}}$ | 2000 | Num. of EC3 model neurons |
| $\mu_{\text{CA3}}$ | $2/N_{\text{CA3}}$ | Mean weights CA3 conns. |
| $\sigma_{\text{CA3}}$ | $\mu_{\text{CA3}}$ | Std. dev. weight CA3 conns. |
| $W_{\text{sum}}^{\text{CA3}}$ | 2 | Initial input sum CA3 conns. |
| $p^{\text{EC3}}$ | 0.05 | Conn. prob. EC3 |
| $w^{\text{EC3}}$ | 0.2 | Conn. weight EC3 |
| $p^{\text{rec}}$ | 0.5 | Conn. prob. recurrent |
| $w^{\text{rec}}$ | 0.005 | Conn. weight recurrent |

**Table 4:** Parameters specifying CA3 model neurons.

| CA3 parameters |  |  |
| --- | --- | --- |
| Parameter | Value | Meaning |
| $r_0^{\text{CA3}}$ | 30 Hz | Maximal CA3 firing rate |
| $\sigma_{\text{PF}}$ | 15 cm | CA3 place field width |

**Table 5:** Parameters specifying EC3 model neurons.

| EC3 parameters |  |  |
| --- | --- | --- |
| Parameter | Value | Meaning |
| $\lambda_{\text{ON} \rightarrow \text{OFF}}$ | 0.6 Hz | Transition rate. ON to OFF |
| $\lambda_{\text{OFF} \rightarrow \text{ON}}$ | 0.04 Hz | Transition rate. OFF to ON |
| $\lambda_{\text{OFF} \rightarrow \text{ON}}^{\text{reward}}$ | 0.16 Hz | Transition rate. OFF to ON<br>around reward site |

For most numerical experiments, the transition probabilities are constant. Only in the case of a reward modulation, *p*_OFF→ON_ are adapted for all model neurons for 1.5 s around the reward zone.

## 9.4 Simulation Experiments

In each experiment, a simulated agent runs multiple laps around a circular track of length 300 cm with a constant velocity *v*. All results in the main part of the study are obtained with *v* = 20 cm*/*s. In the supplement, we systematically vary the velocity to investigate its influence on the size of CA1 model neuron place field widths (Figure S3). At the beginning of a simulation, all dynamic variables (except EC3 model neuron activity) are initialized with 0. EC3 neural activity is initialized with a sample of the stationary distribution of the underlying Markov chain. For Figure 2 and Figure 4, before recording the simulated quantities, the agent runs two laps around the track without being able to trigger plasticity events so that the dynamic quantities can attain values consistent with the simulation. Precise details on the number of runs and repetitions across random seeds are provided below.

At the beginning of the simulations, the spike count threshold *n*_low_ is reduced by 1 for the first 5 laps and *w*^EC3^ is tripled, representing a novelty modulation. In repeated exposure to environments this reduction is not applied.

The single neuron and network parameters remain constant throughout the experiments unless explicitly stated otherwise.

The simulation time step is 1 ms.

## 10 Simulation Details and Data Analyses

All simulations and analyses were performed with customized `python3` code written by ACK with `numpy` version 2.4.6 and scipy version 1.18.0

Analysis of the activity of CA1 model neurons was conducted on the firing rates. We calculated the average activity 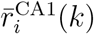 on 2 cm spatial bins on which all subsequent analyses were based. Here, *i* denotes the neuron index and *k* the index of the bin during one lap. We discarded neurons from place field analyses if the peak firing rate 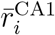 was below 8 Hz or if the firing rate showed two peaks during one lap (assessed by `scipy.signal.find_peaks` with `prominence` set to 2 and `distance` set to 30). The place field center was assessed by fitting a offset Gaussian function to the firing rate using `scipy.optimize.curve_fit`. We computed correlations using `scipy.stats.pearsonr`.

For Figure 2 the agent ran 90 laps on the track. The circular distances between PPs and place fields were computed using the the location of a PP at a given lap and the place field center of the activity on the previous lap. We computed the median of all valid PP-place field distances. Figure 2D shows the meand and standard error of the mean for 5 independent realization of the experiment. For Figure 2E we simulated individual neurons without recurrent inhibitory connections. We induced a PP at the center of the track (150 cm) amd formally set *n*_high_ to infinity to prevent activity induced closing of the *G*(*t*).

For Figure 3 the agent ran 120 laps on the track.

For Figure 4 the agent ran 60 on the track. The reward locations was at 150 cm. 1.5 s around the reward location (i.e. from 130 cm to 160 cm) the EC3 transition probabilities were adapted as described in Table 5, inspired by Grienberger and Magee [13]. For Figure 4C and D we calculated the population means of 5 independent runs, the displayed data shows the mean and standard deviation of the resultant data. For Figure 4E we calculated the correlation between the population-summed CA1 activity and the population summed EC3 activity. We discarded the first 10 laps and computed the mean correlations. We display the distribution of these means computed over 5 independent simulations.

For Figure 5 the agent ran 80 laps independent of how many laps were spent in environment 2. Since the total number of CA3 inputs (but not the number of “active” CA3 inputs in a given environment) was doubled so was the summed CA3 input for the normalization. We systematically increased the time spent in environment 2 in steps of 4 from 0 to (including) 36. For each lap, we computed population vector correlations on the spatially binned data, and then averaged over the obtained correlations of a lap. Correlations were calculated relative to the reference lap 20 (last lap of traversing environment 1). Finally, we assessed variability with the mean and standard deviation of 5 independent simulations. We conducted a weight similarity analysis based on the weights at the end of a lap. We computed the normalized cosine similarity of the weight vector from CA3 to each CA1 neuron between the weights of a given lap and the weights at lap 20. Since all the weights are positive by definition, we subtracted a baseline value *b* from the results and subsequently divided by 1−*b*. We determined the baseline value by calculating the average normalized cosine similarity of the initial weights from two independent instantiations. Again, variability was assessed over 5 random seeds.

For Figure 6 we simulated 500 CA1 neurons without recurrent inhibitory connections. At lap 1 on day 1 we induced a PP at the center of the track (150 cm). One day lasted 10 laps. The exclusion criterion was set to 10 Hz. At the beginning of each subsequent day the weight of EC3 input was tripled. After the first PP of a cell on a given day the value was reset to its default value (see Table 3). At the end of each day the learned weights from CA3 were reduced according to

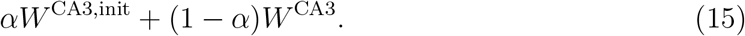

Here, *α* quantifies the strength of the residual trace. After computing the convex combination the weights were normalized as applied at the initialization of the simulations. We varied *α* in steps of 0.125 from 0 (full retention) to 1 (full forgetting). Weak, Moderate, and Strong correspond to *α* = 0.125, 0.375, 0.875 respectively. We determined the location of the PP on a next day to its place field on the previous day from the last measured place field center on the previous day. PP-place field center distance was computed as mentioned above. The reinstantiation of a cell was successful if between a place field center on a given and the previous day was smaller than 30 cm. If there was no PP on a given day this day was skipped. If the conditions were met the cell “survived” this day. For the analysis of the data by Vaidya et al. [33] we used the first PP on a next day on compared them to the last measured place fields on the previous days. In contrast to their analysis, we tracked neurons for which place field centers on subsequent days were not more than 30 cm apart. We grouped data across animals and contexts. Decay strength is defined by grouping the experimental data in disjoint equally sized subsets where, given *N* cells that survived a number of days and *k* groups, the cell with the weakest decay are the *N/k* (rounded if necessary) cells that had the strongest residual activity on the next day assessed by the average fluorescence *±*5 spatial bins around the place field center on the previous day.

## 11 AI Disclosure

Claude Code with Opus 5 was used to understand the source code accompanying the experimental data and to validate the data loading routines used for the reanalysis of the experimental data. Additionally, Claude Code with Opus 5 was used to audit the simulation and analysis code against the descriptions in this manuscript. Consistency was ensured by the authors.

All research questions were conceived, all model development, analysis design, and interpretation of the results were carried out by the authors.

## S1 Supplementary Material

**Figure S1:**
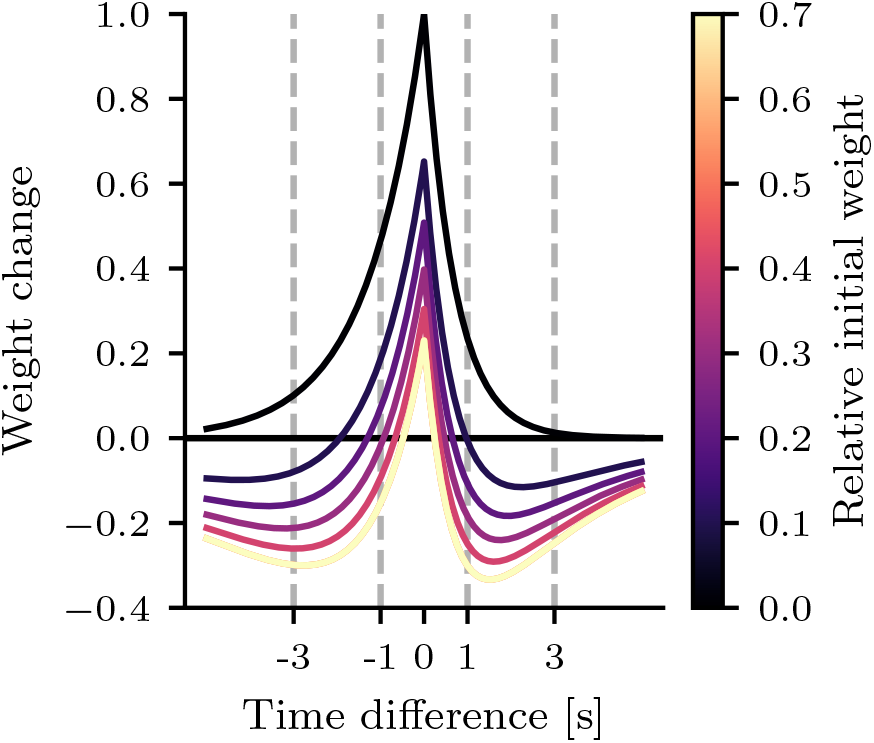
BTSP kernel. The weight change depends on the time difference between the PP and the pre-synaptic activity and the relative size of the initial weight (with respect to the maximal weight).

**Figure S2:**
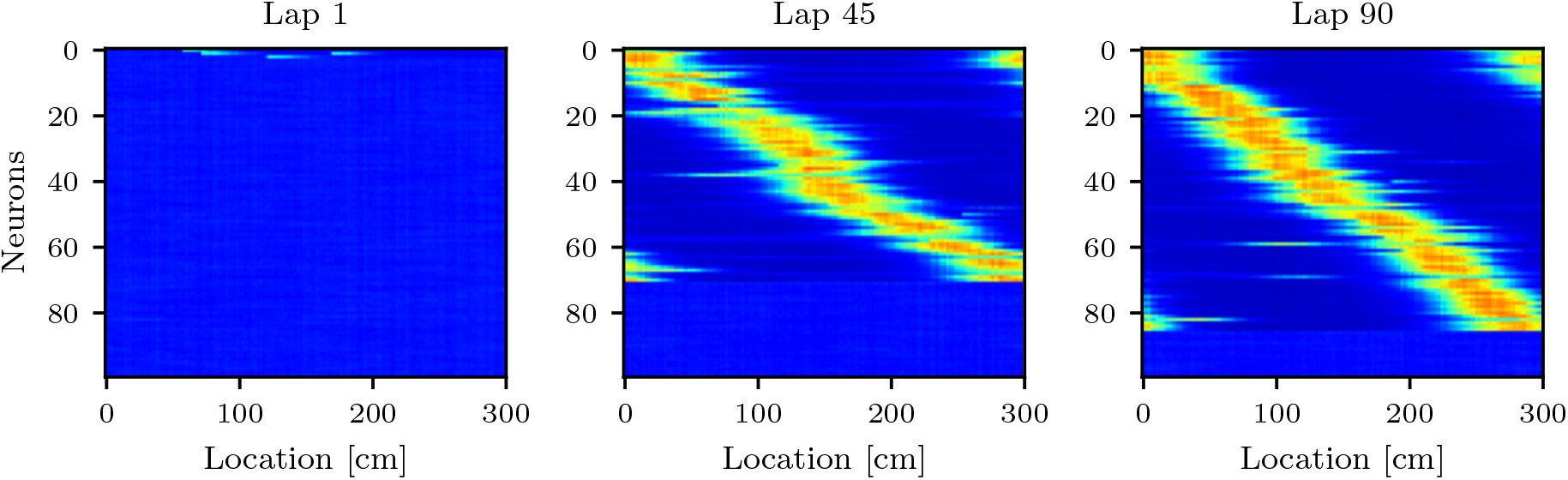
Place field formation without novelty modulation. All other parameters as for Figure 2 (EC3 strength clamped to initial value, no change to default *n*_low_). Sorting according to the respective lap. Left, Lap 1; Center, Lap 45; Right, Lap 90.

**Figure S3:**
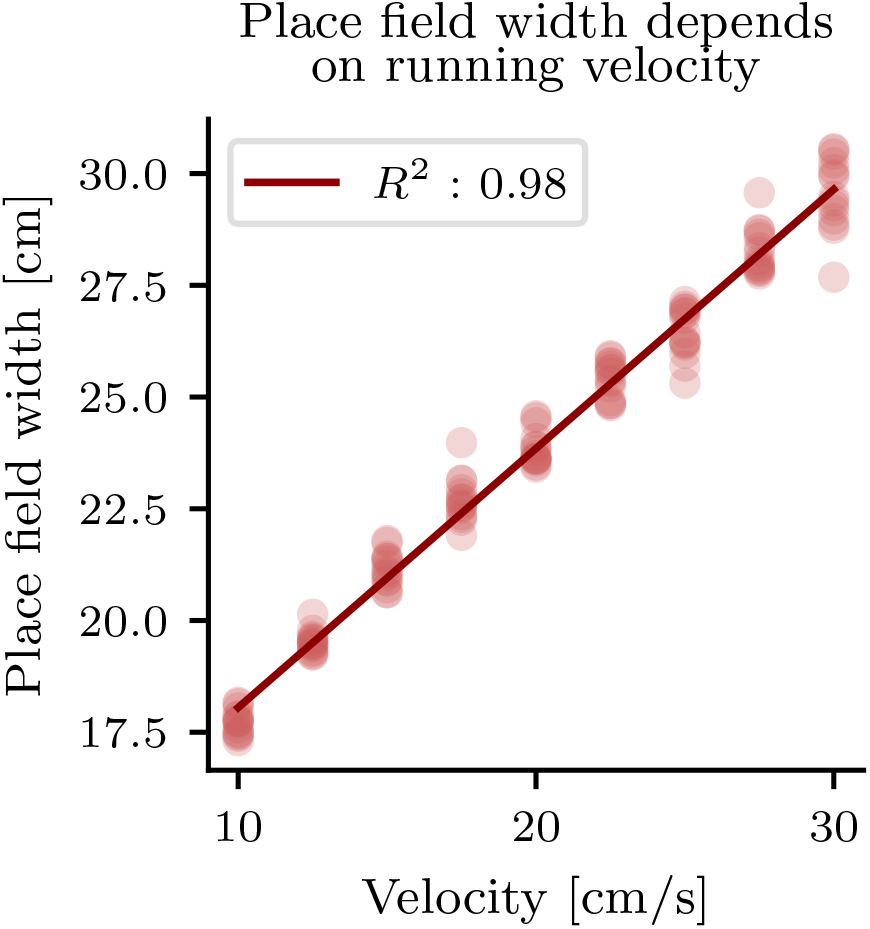
Place field size dependence on running speed. For each velocity, we ran 15 independent single neuron simulations, place fields were induced at the center of the track. Place fields sizes were determined by fitting an offset Gaussian function. The place field width corresponds to the standard deviation. We see a linear relationship between the velocity and the place field width (Pearson correlation).

**Figure S4:**
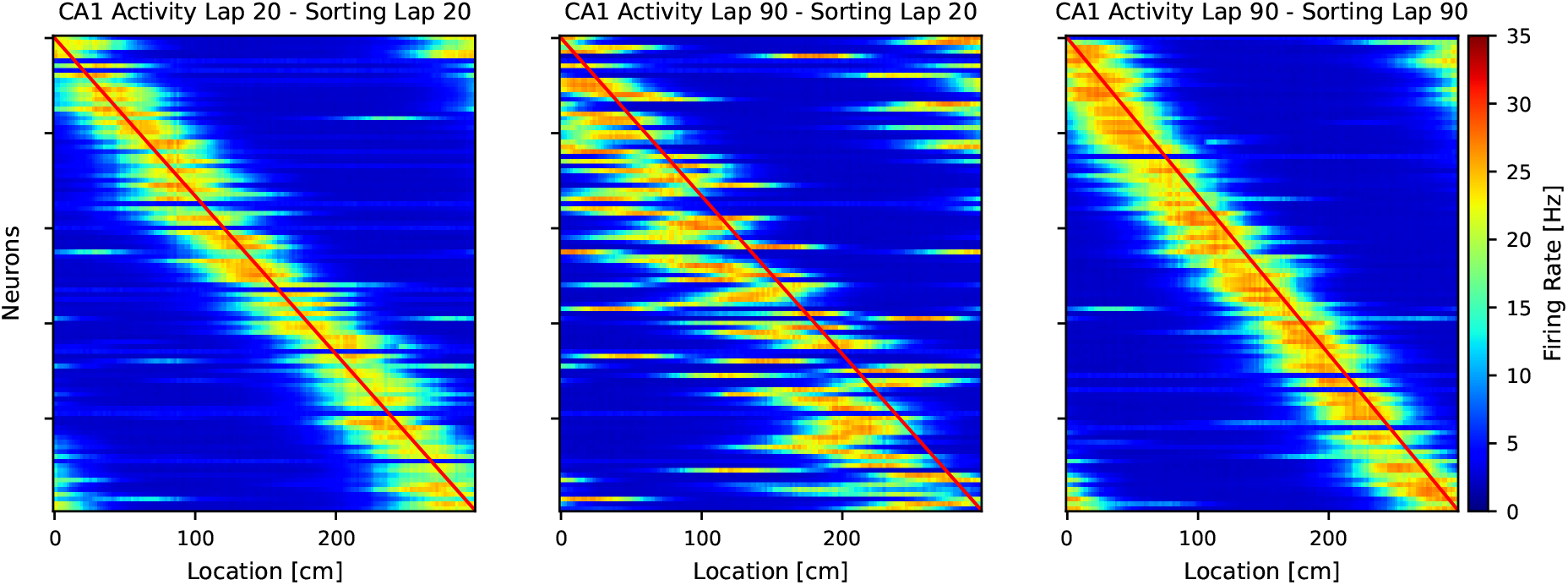
Gradual place field drift. CA1 model activity at different laps. Left: CA1 model activity at lap 20 sorted according to lap 20. Center: CA1 model activity at lap 60 sorted according to lap 20. Right: CA1 model activity at lap 60 sorted according to lap 60. In the center plot, we see, relative to lap 20, noisy drift of the place fields opposite to the running direction.

**Figure S5:**
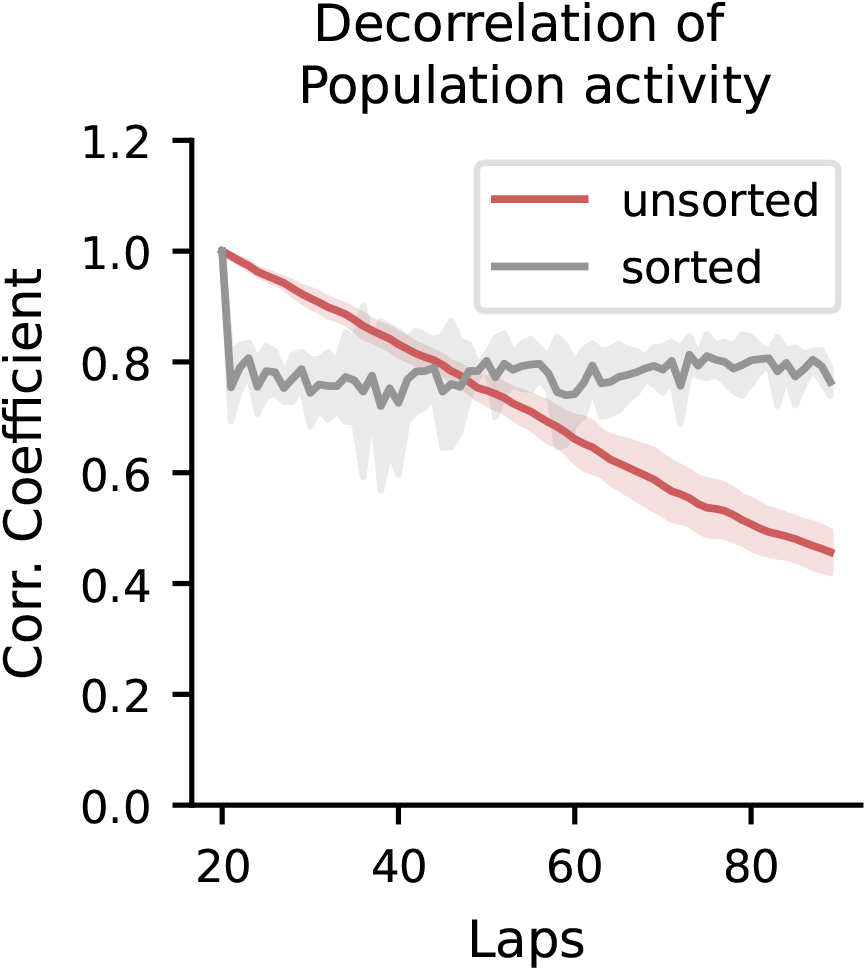
Population correlation between laps. Correlation of between population activity at lap 20 with population activity at later lap averaged over 5 random seed. Shaded area indicates the standard deviation. Activity is the same as used for the analysis in Figure 2. Correlating sorted population activity remains constant at a high level, indicating the same quality of the place field map irrespective of the lap. Comparing non-sorted correlations shows a linear decorrelation with lap, as observed experimentally.

**Figure S6:**
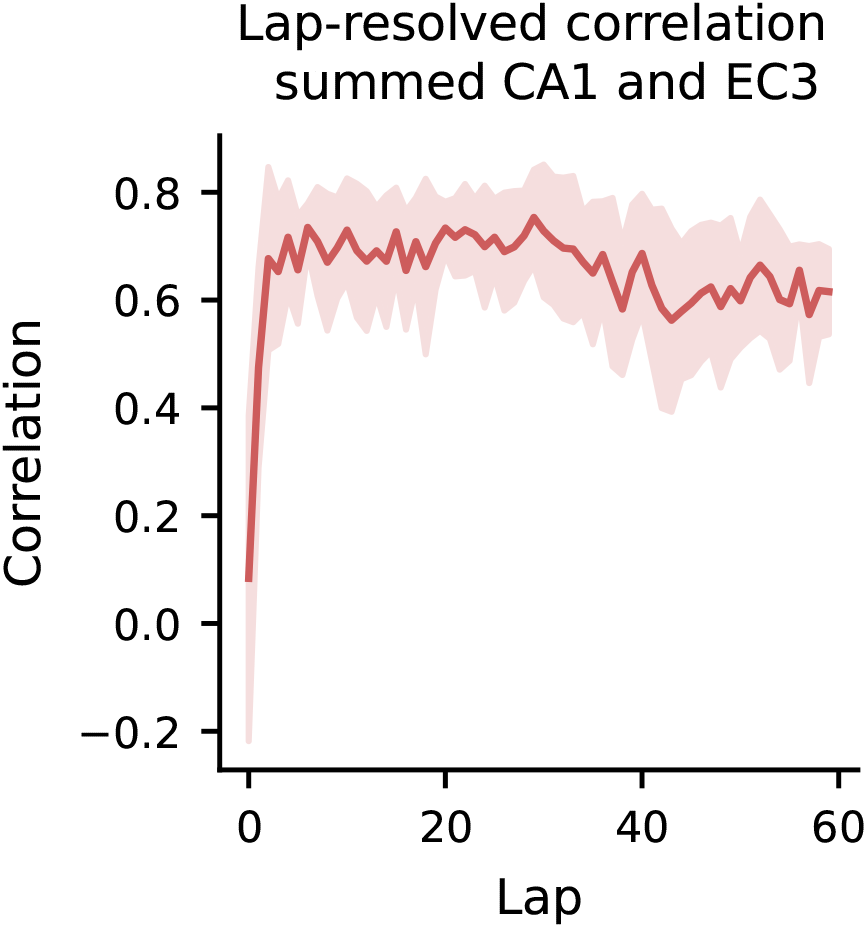
Lap-resolved correlation between summed CA1 and EC3 activity. The correlation increases after a few laps and remains at high level afterwards, as observed experimentally.

**Figure S7:**
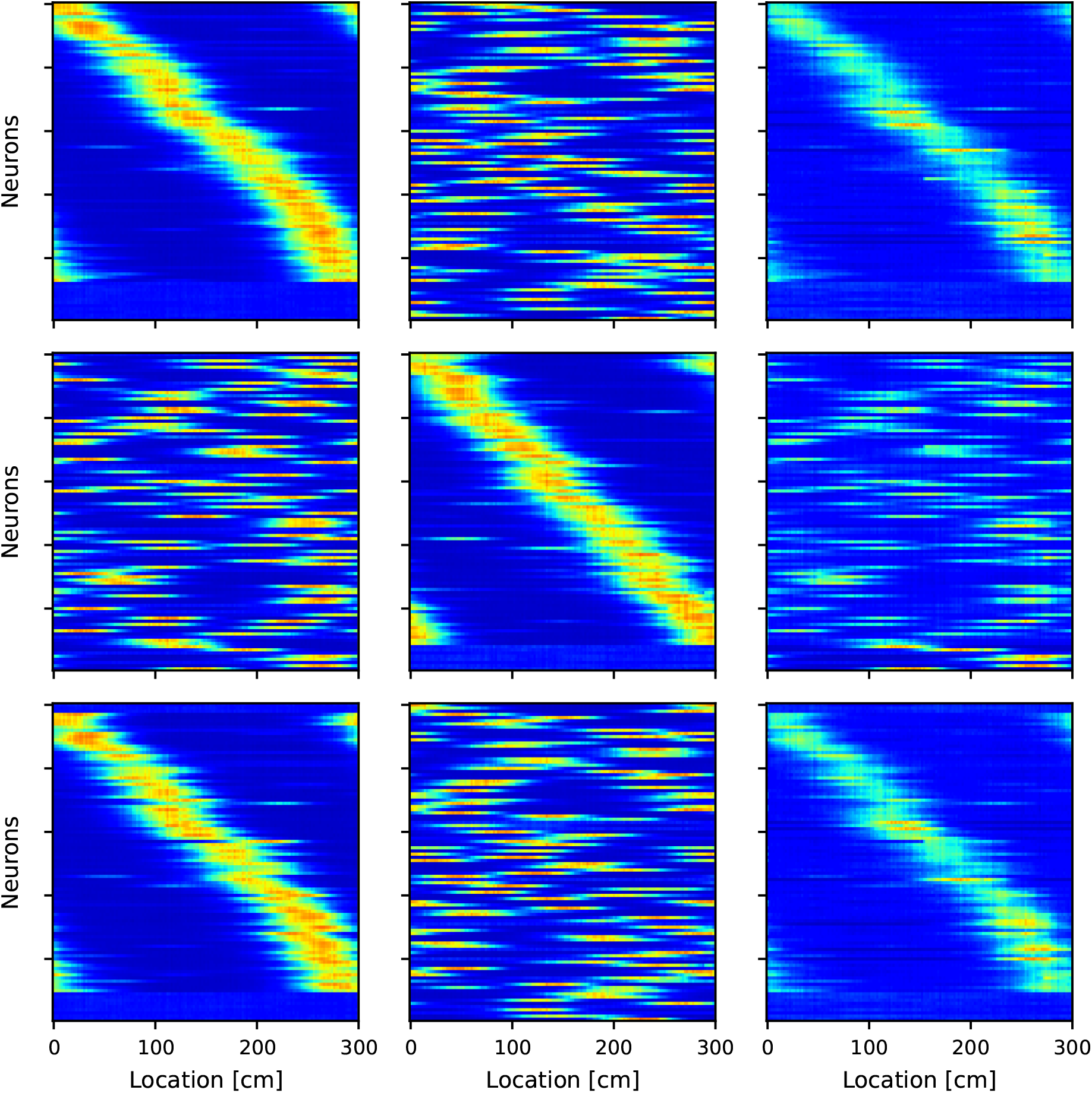
Environment dependent representations. Full cross sorting of activity at lap 20 (last lap in environment 1, first column), activity at lap 44 (last lap in environment 2 after spending 24 laps there, second column), and lap 45 (first lap after reentering environment 1). Activity is sorted according to lap 20 (first row), lap 44 (second row), and lap 45 (third row). The representation formed in environment 2 differs starkly from the representations of environment 1.

**Figure S8:**
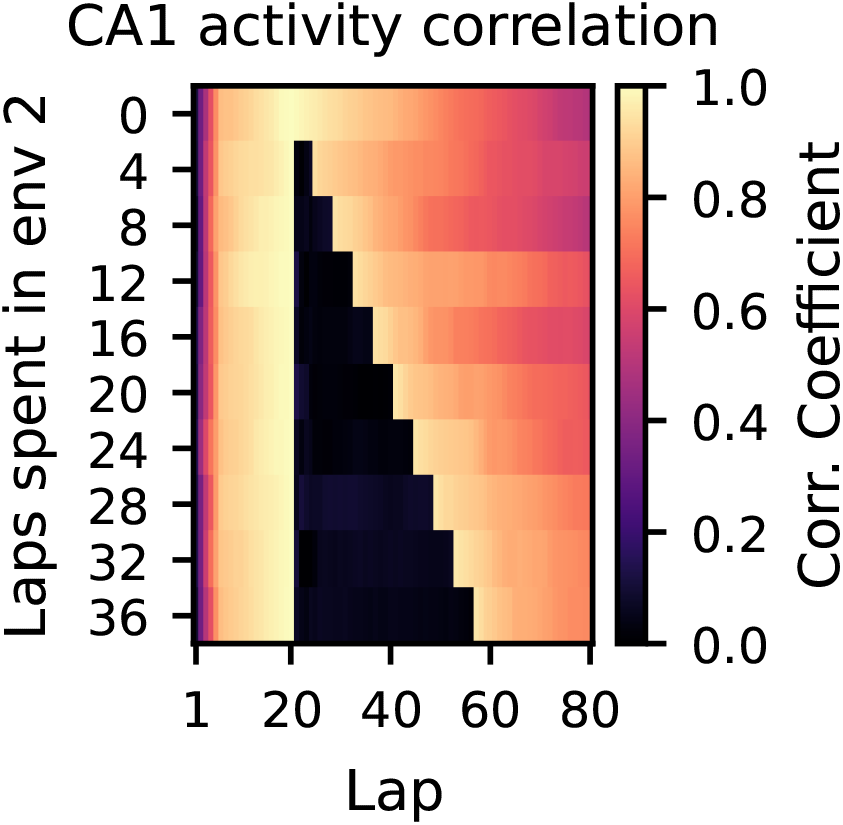
Average CA1 population correlation across laps. Comparison between given lap and lap 20 depending on the number of laps spent in environment 2. The representation formed in environment 2 is fully decorrelated from the representation formed in environment 1. After re-entering environment 1, the representation of environment 1 is immediately re-instantiated and then continues to gradually drift.

**Figure S9:**
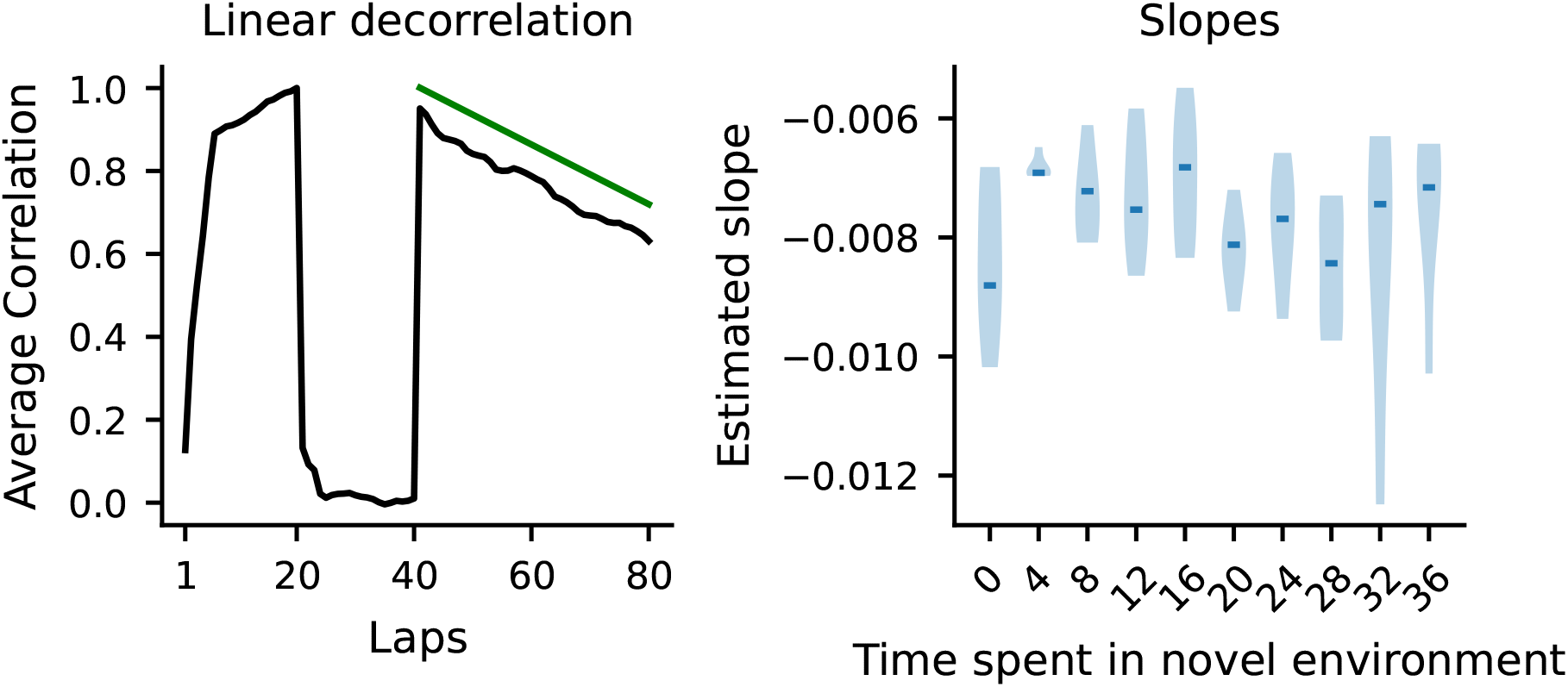
Decorrelation speed dependent on time spent in novel environment. Left: Example average CA1 population correlation of given lap and lap 20. 20 laps spent in environment 2. Green line results from linear regression of the decorrelation after re-entering environment 1 (offset for illustration purpose). Right: Distribution of slopes of decorrelation.

